# Neural division of labor: the gastropod *Berghia* defends against attack using its PNS to retaliate and its CNS to erect a defensive screen

**DOI:** 10.1101/2023.07.29.551068

**Authors:** Jeffrey W. Brown, Ondine H. Berg, Anastasiya Boutko, Cody Stoerck, Margaret A. Boersma, William N. Frost

## Abstract

Relatively little is known about how the peripheral nervous system (PNS) contributes to the patterning of behavior, in which its role transcends the simple execution of central motor commands or mediation of reflexes. We sought to draw inferences to this end in the aeolid nudibranch *Berghia stephanieae*, which generates a rapid, dramatic defense behavior, “bristling.” This behavior involves the coordinated movement of cerata, dozens of venomous appendages emerging from the animal’s mantle. Our investigations revealed that bristling constitutes a stereotyped but non-reflexive two-stage behavior: an initial adduction of proximate cerata to sting the offending stimulus (Stage 1), followed by a coordinated radial extension of remaining cerata to create a pincushion-like defensive screen around the animal (Stage 2). In decerebrated specimens, Stage 1 bristling was preserved, while Stage 2 bristling was replaced by slower, uncoordinated, and ultimately maladaptive ceratal movements. We conclude from these observations that 1) the PNS and central nervous system (CNS) mediate Stages 1 and 2 of bristling, respectively; 2) the behavior propagates through the body utilizing both peripheral- and central-origin nerve networks that support different signaling kinetics; and 3) the former network inhibits the latter in the body region being stimulated. These findings extend our understanding of the PNS’s computational capacity and provide insight into a neuroethological scheme that may generalize across cephalized animals, in which the CNS and PNS both independently and interactively pattern different aspects of non-reflexive behavior.

## Introduction

Many seminal breakthroughs in neuroethology have exploited the relatively simple central nervous systems (CNS) belonging to a cadre of model invertebrate species (e.g., *C. elegans*, *Drosophila, Hirudo, Aplysia*) to derive fundamental causative relationships between brain and behavior that often generalize across animal phylogenies [1-4]. Do peripheral nervous systems (PNS), whose more mundane roles as participants in local reflexes and downstream effectors of central motor outputs are more thoroughly understood, likewise possess sufficient computational power to cooperatively or independently generate behavior? The answer to this question has remained largely elusive owing to the comparatively scant body of literature characterizing the peripheral neuroanatomy of many canonical model species; this is a reflection of both the typically small size and staggering number of peripheral neurons, and the resulting difficulty in designing meaningful neuroethological investigations into the peripheral correlates of behavior. While the octopus, for example, has proved a uniquely profitable model species in which to study how complex behavior arises from sensorimotor integration across confederated but not strictly hierarchical nervous systems [5-8], the size of the cephalopod nervous system (on the order of 500 million neurons) and commensurate physical complexity of cephalopod movement make these animals relatively challenging models in which to study the neural bases of behavior at granular levels of association.

Elsewhere in the phylum Mollusca, the gastropods, which possess between 5,000 and 25,000 often large, uniquely identifiable central neurons [9] and unsegmented bodies with specialized appendages, occupy a happy medium within the pantheon of animal species commonly used in neuroethological research: with respect to anatomical, neural, and behavioral complexity, gastropods sit between the 302-neuron-strong brain and streamlined morphology of *C*. *elegans* and the more elaborate brains and body designs of cephalopods, arthropods, and vertebrates. Research in the CNS of gastropods has elucidated fundamental, broadly conserved principles of neurophysiology, such as long-term potentiation and the cellular and molecular bases of both nonassociative and associative learning [10-13], central pattern generation [14], neuromodulation [15-18], and neuroeconomics [19-22]. Although investigations in these animals have additionally revealed specific contributions made by the PNS to several reflexes [23-26], the manners in which the CNS and PNS interact to generate more dynamic behaviors in these animals remains unexplored.

In seeking to utilize a gastropod model to this end, we focused our attention on *Berghia stephanieae* [27], an aeolid nudibranch indigenous to the Caribbean Sea and a commercially available “anemone cleaner” popular with aquarists [28]. A particularly flamboyant, uncharacteristically rapid behavior exhibited by this animal is the subsecond, coordinated, targeted convergence and/or elevation of the cerata, appendages emerging from the animal’s mantle, in response to a noxious stimulus (Fig. 1A; Video S1; see Fig. 1B for a diagram of *Berghia*’s relevant anatomy). This defensive behavior, dubbed “bristling” [29], has been documented in other aeolid nudibranchs in response to both assault from predators and during the animals’ interaction with their cnidarian prey [29-32]. Like other aeolids, *Berghia* co-opts nematocysts, specialized cells that explode to launch a toxin-laden tubule into predators, through the consumption of its principal anemone prey, *Exaiptasia diaphana*, which produces these structures endogenously and sequesters them in cnidosacs, which are situated at the tips of the cerata [27, 30, 33-35].

**Figure 1.**
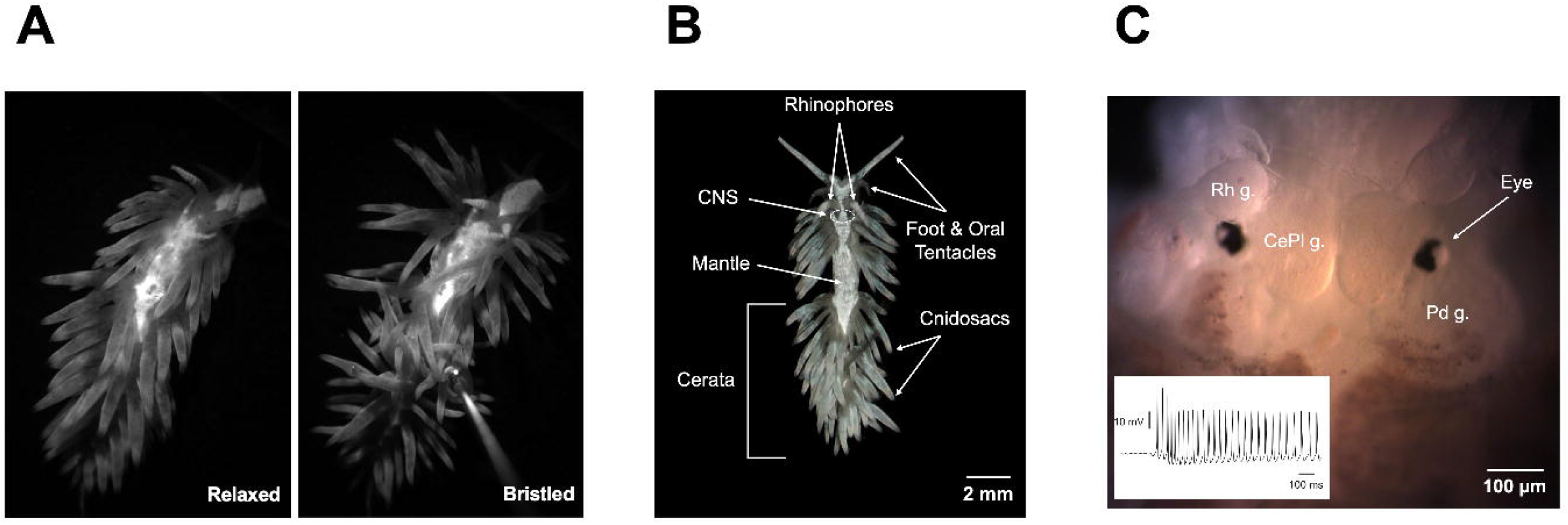
Tactile-elicited bristling in and the associated general anatomy and central nervous system of *Berghia stephanieae.* (A) Bristling arises in under a second following stimulation and involves the rapid deployment of cerata, the dozens of finger-like appendages emerging from the animal’s mantle, to either converge on and sting the offending stimulus or to elevate and/or extend outwardly around the animal’s body. The left and right panels respectively show relaxed (pre-stimulatory) and fully “bristled” states. See also Video S1. (B) The general anatomy and (C) central nervous system (CNS) of *Berghia stephanieae*. The inset in Panel C features a sample sharp-electrode recording from a neuron located in the cerebropleural ganglion (CePl g.) of a whole-animal preparation. Rh g., rhinophore ganglion; Pd g., pedal ganglion.

That *Berghia* seemed capable during our preliminary observations of exercising selective motor control over individual ceratal clusters during bristling invited our speculation as to whether the patterning of this behavior may be distributed across the animal’s CNS (Fig. 1C) and PNS, a supposition grounded in existing literature demonstrating the involvement of both central and peripheral substrates in moving or automizing cerata in other gastropod species [32, 36, 37]. In the present study, we characterize the ceratal kinematics inherent to bristling in both intact and decerebrated *Berghia* through the application of aversive stimuli at several dorsal body loci and analyze how these kinematics change following the removal of the CNS. From this analysis, we draw general inferences about how an animal’s CNS and PNS may interactively generate different aspects of a directed, non-reflexive behavior and moreover posit a rudimentary anatomical model of the neural infrastructure underlying it.

## Results

### Bristling involved a two-stage response: inward targeting of the stimulus by local cerata and outward radiation of other cerata to erect a protective screen around the body

To facilitate high-magnification, high-speed resolution of bristling kinematics, we immobilized *Berghia* specimens (N=20) within the field of view of a Zeiss Stemi SV-11 stereomicroscope (Carl Zeiss Microscopy, White Plains, NY) using a customized vacuum apparatus (Fig. S1A). Videos of bristling animals were acquired using a C-mounted Hamamatsu ORCA-Flash4.0 V3 sCMOS camera (Bridgewater, NJ). Specimens were stimulated through the application of a 3.61-gauge von Frey hair (VFH; Stoelting Company, Wood Dale, IL) to one of three dorsal body loci: the head, midbody, and caudal targeting sites; cerata were classified based on their proximities to these three stimulation loci (Fig. S1B).

Bristling consisted of a sequence of two complementary ceratal behaviors: an initial, rapid targeting of the offending stimulus by the nearest cerata, dubbed Stage 1 bristling, followed by a coordinated radial extension of most distal cerata along the length of the animal’s body that amplified the effective body volume by creating a defensive screen, or Stage 2 bristling. In freely locomoting animals, bristling was accompanied by concomitant defenses, including body shortening, head withdrawal, and avoidance turning, whose magnitudes and induction rates varied as a function of where specimens were stimulated (Figs. S2 and S3; Videos S2 and S3). The general two-step bristling sequence was observed consistently across the three body regions to which the VFH was applied (Fig. 2; ceratal movement is best discerned in Videos S4-S6). Although application of the VFH to the three selected loci along the mantle generated a modest local depression at the point of contact, the angles to which cerata participating in either stage of bristling rotated away from their equilibrium positions were consistently too large and the rotational directions generally too incongruent with the downward force applied by the VFH for ceratal pivoting to be attributable to mere mechanical coupling with the mantle.

**Figure 2.**
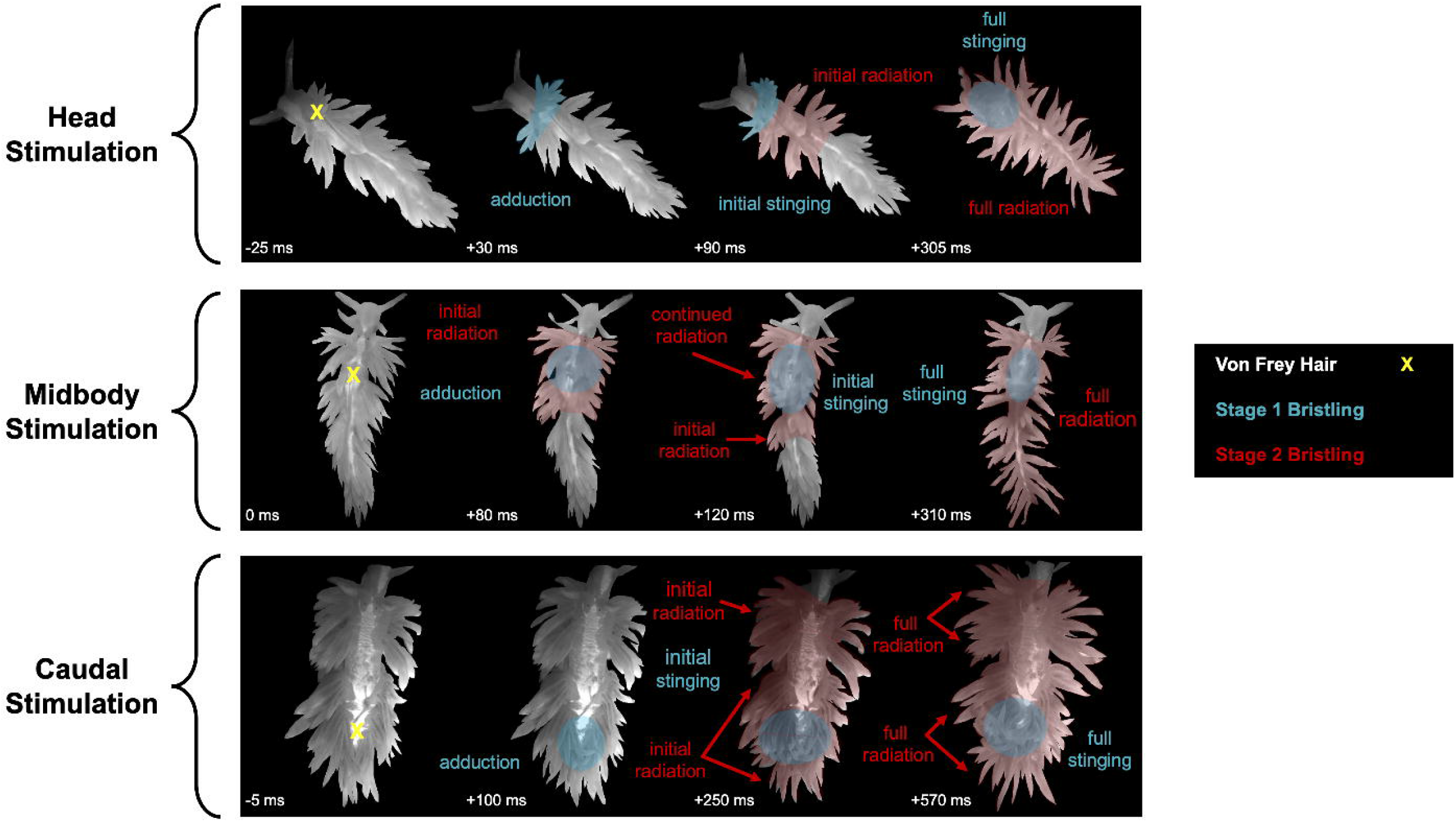
Bristling consisted of a two-stage response, wherein local cerata pivoted inward to target the stimulus while other cerata radiated outward to form a protective screen around the animal. Representative sequences of bristling in response to punctate tactile stimulation of the head, midbody, and caudal regions in vacuum-immobilized *Berghia* with a 3.61-gauge von Frey hair (VFH) are shown. Stage 1 of the bristling response manifested in the cerata closest to the VFH stimulus (denoted by the “X”) and involved adduction or centripetal pivoting towards and ultimately direct targeting of the stimulus by these appendages; this stage of bristling can be visualized more clearly in Videos S4-6. A second stage of bristling commenced following the initiation of Stage 1: this phase of the behavior was characterized by a coordinated, outward radiation of cerata found more distal to the stimulus. Dorsally situated cerata participating in Stage 2 adducted and elevated above the mantle, while those cerata originating along the lateral aspects of the body pivoted perpendicularly to it. Ceratal movement in Stage 2 generated a pincushion-like screen around the body that extended the animal’s defensive radius. Time indices in each subpanel are reported in relation to stimulation.

Stage 1 bristling involved a directed adduction and wrapping of the nearest cerata around the stimulus, with other nearby cerata anterior or posterior to the VFH pivoting centripetally toward the stimulus without directly contacting it. While vigorous targeting of the stimulus by sufficiently close cerata was observed in all cases of head and caudal stimulation and in a majority of midbody stimulations, milder wrapping around the VFH was sometimes elicited in the midbody (N=8 of 19), possibly reflecting a lower density of cerata within reach of this region relative to more anterior or posterior loci.

Subsequent to the directed ceratal retaliation marking Stage 1 bristling, a second stage of bristling was marked by a coordinated, outward radiation of cerata found more distal to the stimulus, with these cerata typically dispersing from one another to create a voluminous, pincushion-like protective screen around the body. For the dorsally situated cerata recruited during Stage 2 bristling, outward radiation entailed elevation above the mantle and adduction, while cerata originating along the lateral aspect of the body pivoted perpendicularly thereto, extending the animal’s defensive radius within the horizontal anatomical plane. Regardless of their points of origin, all cerata participating in Stage 2 moreover pivoted anteriorly while being recruited into the behavior.

### Temporally distinct sequences of ceratal recruitment during bristling depended on stimulation locus and revealed different modes of bristling propagation along the body

To determine the sequence of ceratal recruitment characteristic of bristling elicited through tactile stimulation to the head, midbody, and caudal loci, we measured the post-stimulus latencies with which cerata in each of these three body regions were initially recruited into the bristling response; additional post-stimulus latencies to full-body ceratal recruitment (i.e., the time required for all cerata to respond to stimulation) and to the time at which maximal bristling magnitude was achieved were compared across stimulation regions (Fig. S4), as were bristling magnitudes themselves (Fig. S5). Across all trials, cerata originating in the region being stimulated were the first to respond: this was consistent with the observation that the cerata most proximate to the VFH directly targeted it during Stage 1 bristling. Beyond this initial bristling stage, the order of subsequent ceratal recruitment into Stage 2 bristling varied depending on the body locus stimulated. In trials featuring head stimulation (N=15), cerata were consistently recruited in a head-midbody-caudal sequence (Fig. 3, top row), with average post-stimulus latencies of 17.3 ± 3.6, 54.0 ± 9.4, and 141.3 ± 23.9 ms, respectively; this indicated a posteriorly propagating wave of bristling initiation in which cerata in one region responded with distinct latencies relative to those originating in the other two regions [one-way RM ANOVA, *F*(2,14)=70.64, *p*<0.0001; all pairwise Tukey’s tests, *p*<0.0001]. Cerata in all three body regions responding to midbody stimulation (N=19) similarly activated with distinct latencies but in a midbody-head-caudal sequence in all instances (average latencies: midbody, 33.7 ± 4.9 ms; head, 77.6 ± 10.9 ms; caudal, 212.1 ± 25.6 ms), apparently reflecting the shorter transit distance between the midbody and head than the midbody and caudal region [Fig. 3, middle row; *F*(2,18)=48.96, *p*<0.0001; Tukey’s tests: *p*=0.0002 for head vs. midbody, *p*<0.0001 for head vs. caudal and midbody vs. caudal].

**Figure 3.**
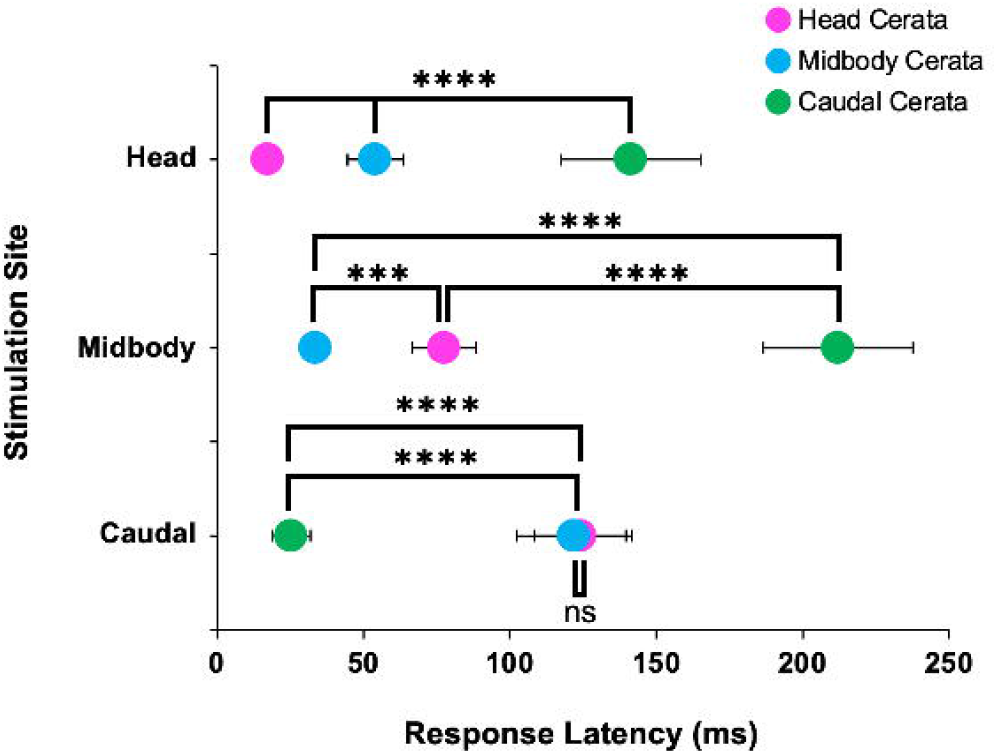
Sequences of ceratal recruitment during bristling depended on tactile stimulation locus and revealed distinct modes of bristling propagation. Cerata in the body region being stimulated, most of which participated in Stage 1 bristling, were consistently the first appendages recruited into the bristling response and thus exhibited the shortest response latencies (depicted here as mean ± standard error). During stimulation of the head (N=15) and midbody loci (N=19), the post-stimulus response latencies of cerata originating in each body region were temporally distinct and increased as a function of increasing distance between the stimulus site and body region in question. By contrast, when the caudal body was stimulated (N=15), there was no significant difference in the recruitment latency between midbody and head cerata, despite a significant main effect across body regions. The displayed statistics reflect Tukey’s post-hoc tests run in conjunction with one-way ANOVAs. ****, *p*<0.0001, ***, *p*<0.001; ns, not significant.

Although caudal cerata were predictably the first to respond to caudal stimulation [N=15; average caudal latency, 25.3 ± 6.6 ms; *F*(2,14)=55.52, *p*<0.0001; Tukey’s tests, *p*<0.0001 for head vs. caudal and midbody vs. caudal], there was surprisingly no difference between the average latencies with which midbody and head cerata (122.0 ± 19.7 and 124.0 ± 15.5 ms, respectively; *p*=0.597) were recruited into the bristling response (Fig. 3, bottom row); in a plurality of caudal-stimulation trials (N=7 of 15), midbody cerata activated before head cerata, with head cerata preceding midbody cerata in four cases and both ceratal groups responding simultaneously in four other trials. The unexpected finding that bristling induction in the head and midbody regions occurred with statistical simultaneity when the caudal locus was stimulated suggested that the afferent signals ultimately inducing Stage 2 bristling were either 1) integrated in the CNS or 2) traveled along distinct afferent pathways with different propagation velocities and were integrated locally in the midbody or cephalic PNS; were these signals integrated peripherally along the same pathway, midbody cerata should have consistently responded before head cerata based on the former appendages’ proximity to the caudal stimulation locus. To assess these two mechanistic possibilities, we pursued studies in decerebrated animals.

### Decerebration left Stage 1 bristling intact while eliminating Stage 2 bristling, respectively implicating the PNS and CNS in these phases of the behavior

Aspects of the stimulus-elicited bristling characterized in intact animals, most notably Stage 1, persisted in decerebrated specimens (N=10), in whom the CNS had been excised 24 hours prior and the dorsal incision enabling CNS excision allowed to seal (Fig. 4; Video S7). Specimens lacking a CNS typically exhibited a high degree of uncoordinated, spontaneous ceratal movement throughout the body that persisted in the absence of stimulation (Video S8), possibly reflecting the deprivation of central tonic inhibition prevailing in intact animals not under threat, whose cerata remained relatively motionless. Nevertheless, all elements of Stage 1 bristling were observed in decerebrated animals, albeit with a greater variability of intensity and frequency; this variability might have owed to either a loss of excitatory central neuromodulation of peripheral motor elements or to the fact that Stage 1 cerata were sometimes already in motion at the time of stimulation. Cerata close to the impinging VFH typically adducted either vigorously (head, N=4 of 10; midbody, N=5 of 10; caudal, N=6 of 10) or minimally (head, N=4 of 10; midbody, N=5 of 10; caudal, N=3 of 10), though there were a minority of both head (N=2 of 10) and caudal trials (N=1 of 10) in which this response was altogether absent. A subset of these proximate cerata in decerebrated specimens furthermore wrapped around and stung the VFH, as in intact animals: this occurred vigorously (head, N=3 of 10; midbody, N=5 of 10; caudal, N=5 of 10), minimally (head, N=2 of 10; midbody, N=3 of 10; caudal, N=3 of 10), or not at all (head, N=5 of 10; midbody, N=2 of 10; caudal, N=2 of 10).

**Figure 4.**
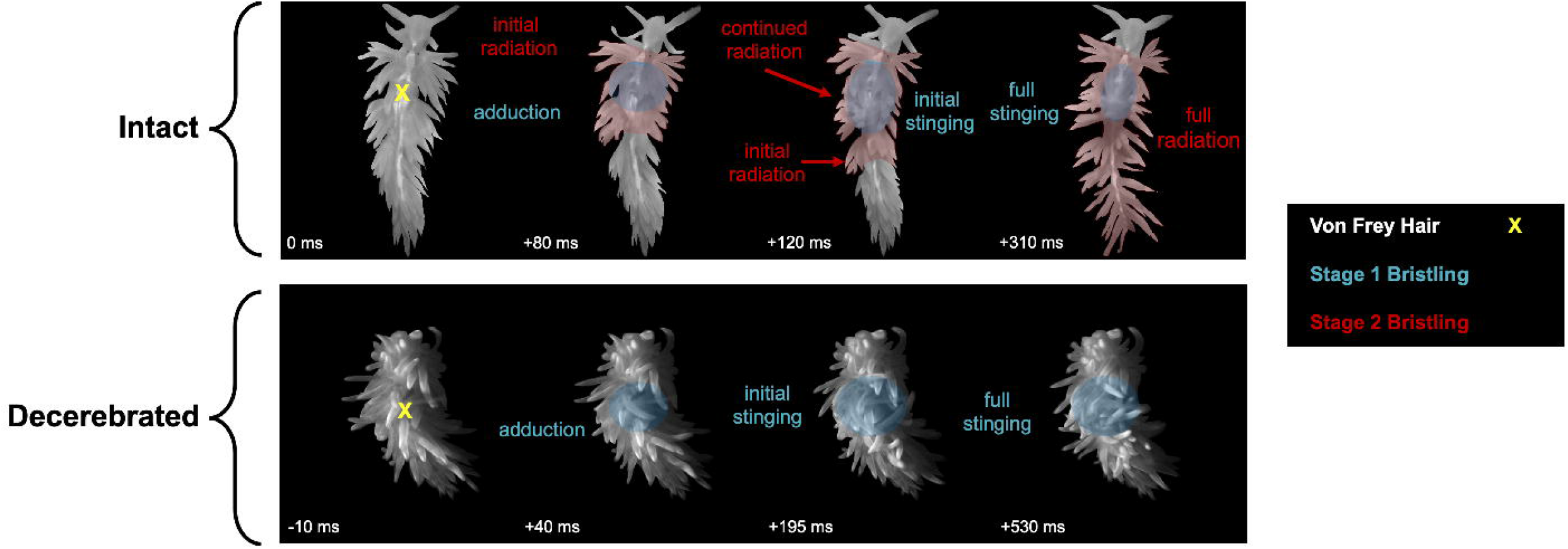
Stage 1 bristling was preserved though sometimes attenuated in decerebrated *Berghia*, while Stage 2 bristling was abolished in response to tactile stimulation. Shown here are representative sequences of bristling in intact and decerebrated *Berghia*, respectively, the latter of which had its CNS removed 24 hours prior to experimentation, responding to tactile stimulation of the midbody. Cerata in decerebrated animals nearest to the impinging von Frey hair adducted or centripetally pivoted to sting the stimulus, mirroring the initiation of Stage 1 bristling in intact animals. The subsequent wrapping of some of these cerata around the VFH occurred in only some trials featuring decerebrated animals, including the specimen shown here, and then with a lower average density of converging cerata than in intact counterparts. Absent from decerebrated animals was the outward radiation and dispersion of non-local cerata intrinsic to Stage 2 bristling as observed in intact animals. Time indices in each subpanel are reported in relation to stimulation. See also Videos S5 and S7.

While cerata outside the radius of those exhibiting the targeted, Stage 1 bristling response in decerebrated animals sometimes moved in response to tactile stimulation as well, they did so in manners markedly inconsistent with Stage 2 bristling as observed in intact animals, with Stage 2 bristling replaced by an uncoordinated, undirected, and ultimately maladaptive mode of ceratal movement. Such cerata in decerebrated specimens lacked the concerted, radiative pivoting and dispersion witnessed in Stage 2 bristling, instead pivoting far more modestly and then either inwardly or in seemingly random directions, as constrained by their ranges of motion. Moreover, whereas all cerata recruited into both stages of bristling in intact animals remained relatively fixed in position for some time following their initial excursion before relaxing, non-local cerata in decerebrated animals frequently exhibited undirected wobbling or, in some cases, more dramatic, oscillatory flailing after their initial pivots, with the magnitude of these movements typically eclipsing the spontaneous ones observed before stimulation. The seemingly unregulated persistence of movement by certain cerata appeared to derive in part from collisions between appendages, wherein one ceras would induce movement in another. Indeed, this ceratal coupling, which was conspicuously absent in intact animals in whom bristling entailed highly coordinated movements across cerata, appeared sometimes to underlie the spread of non-local ceratal responses in decerebrated animals.

Further distinguishing the behavior of non-local cerata in decerebrated specimens was the extent of recruitment along the length of the animal: while Stage 2 bristling in intact animals almost always entailed the recruitment of *all* non-local cerata into the behavior, the distance from which waves of haphazard, maladaptive ceratal movement radiated from the stimulation site varied considerably across decerebrated specimens. Head stimulations sometimes generated waves that propagated down the full length of the body (N=4 of 10), though they more commonly terminated in the midbody region (N=6 of 10). Non-local ceratal movement in response to midbody stimulation was more variable: in some instances, cerata in the head and caudal regions (anterior and posterior to the stimulus, respectively) responded (N=4 of 10), while other trials saw only caudal responses (N=2 of 10) or a lack of non-local ceratal movement (N=4 of 10). When the caudal region was stimulated in decerebrated animals, the ensuing response wave typically terminated in the midbody (N=6 of 10), with further head cerata recruited (N=2 of 10) or a lack of non-local response propagation (N=2 of 10) in other cases. These results collectively suggested that while Stage 1 bristling was fundamentally retained in the absence of the CNS and could therefore be attributed to the PNS, Stage 2 bristling depended on central sequencing and coordination.

Despite the absence of Stage 2 bristling and the abundance of spontaneous ceratal movement in decerebrated *Berghia*, two kinematic measurements could nevertheless be quantified in these specimens and consequently compared to corresponding measurements made in intact animals. First, the retention of Stage 1 bristling in decerebrated specimens enabled us to measure the post-stimulus latency to the onset of bristling (i.e., the initial stimulus-evoked Stage 1 movement observed in proximate cerata). We employed a two-way ANOVA to test whether either stimulus locus and/or CNS status influenced this metric (intact animals undergoing head stimulation, N=15; midbody, N=19; tail, N=15; decerebrated animals undergoing head stimulation, N=6; midbody, N=5; tail, N=4). This analysis revealed no statistically significant interaction between stimulus site and CNS status on Stage 1 bristling onset latency [*F*(2,58)=0.394, *p*=0.677], with simple main effects analyses likewise indicating no significant independent effect of either stimulus locus (*p*=0.219) or CNS status (*p*=0.788) on onset latency (Fig. 5A). The latter result, which implied that the CNS played no role in expediting the onset latency of Stage 1 bristling, further underscored the sufficiency of the PNS in exclusively driving this portion of the behavior.

**Figure 5.**
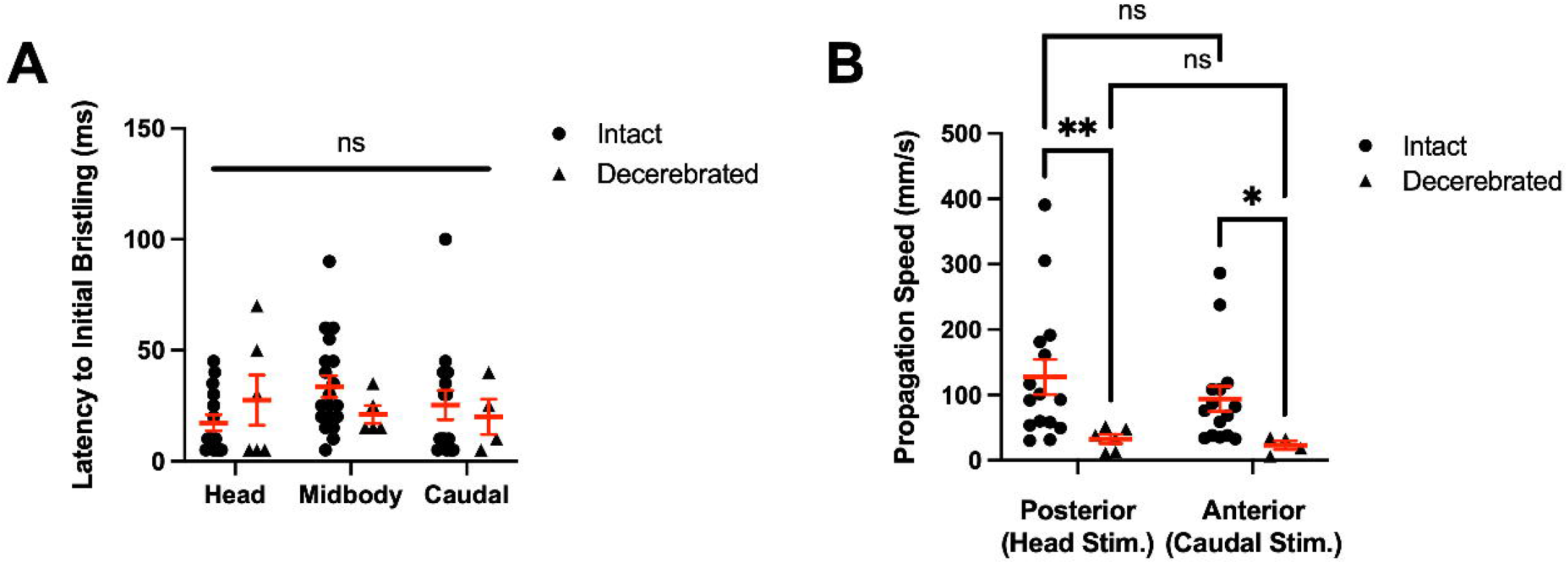
Decerebration significantly slowed the speed at which tactilely elicited waves of ceratal recruitment propagated through the body, while exerting no effect on the post-stimulus latency to the onset of bristling. (A) There were no significant differences in the post-stimulus latencies to bristling onset (i.e., the initial stimulus-evoked ceratal movement, which was consistently observed in proximate cerata participating in Stage 1 bristling) exhibited by way of any interaction between stimulus locus and CNS status, nor through independently exerted effects by each independent variable. (B) While there was neither any significant interaction between the direction of bristling wave propagation (posterior from head stimulation or anterior from caudal stimulation) and CNS status nor a simple main effect of propagation direction on propagation speed, decerebration was shown to significantly slow ceratal wave propagation in both the posterior and anterior directions. Sample sizes for intact specimens subjected to stimulation at each body locus: head, N=15; midbody, N=19; tail, N=15. Sample sizes for decerebrated specimens: head, N=6; midbody, N=5; tail, N=4. Means and standard errors are overlayed on data distributions. The displayed statistics reflect Tukey’s post-hoc tests run in conjunction with two-way ANOVAs. **, *p*<0.01; *, *p*<0.05; ns, not significant.

Secondly, to ascertain whether the rate at which waves of ceratal recruitment propagated through the body was dependent on propagation direction and/or CNS status, we converted both posterior (head-to-caudal) and anterior (caudal-to-head) bristling induction latencies measured in intact animals stimulated at the head and caudal loci, respectively, to propagation speeds by accounting for head-to-caudal distances; (midbody stimulations, in which bristling waves propagated bidirectionally, were not considered in the interest of simplicity). This transformation was implemented to accommodate those decerebrated animals in whom ceratal recruitment waves failed to spread throughout the full body in a manner consistent with Stage 2 bristling but in whom the ceratal recruitment delay between two arbitrary body loci could be nonetheless measured to generate comparable propagation speed measurements. A two-way ANOVA did not disclose either a significant interaction between the effects of propagation direction and CNS status [*F*(1,36)=0.048, *p*=0.873] or an individual effect of propagation direction on propagation speed (*p*=0.241). However, simple main effects analysis indicated a highly significant effect of CNS status on propagation speed (*p*<0.0001), whereby both posterior and anterior ceratal recruitment waves propagated more rapidly in intact (posterior propagation speed, 127.7 ± 26.9 mm/s; anterior propagation speed, 94.0 ± 19.3 mm/s) than in decerebrated animals (posterior, 32.7 ± 7.0 mm/s; anterior, 23.2 ± 6.6 mm/s) (Fig. 5B; Tukey’s tests: *p*=0.0061 for posterior propagation and *p*=0.0107 for anterior propagation). We therefore deduced that the CNS plays a critical role not only in coordinating and sequencing the whole-body bristling response exhibited in Stage 2 of the behavior, but also in expediting it. Table 1 summarizes the mechanistic differences in bristling noted between intact and decerebrated *Berghia*.

**Table 1.** A comparison of mechanistic aspects of bristling between intact and decerebrated *Berghia* specimens. H, head region; M, midbody region; C, caudal region.

|  | Intact | Decerebrated |
| --- | --- | --- |
| **Baseline Ceratal Movement** | Negligible, cerata similarly oriented across body | Significant, random in direction, uncoordinated |
| **Initiation of Bristling (Stage 1)** | Proximal to stimulus; equally rapid |  |
| **Wrapping of Cerata Around Stimulus (Stage 1)** | Typically vigorous<br>(H: N=15 of 15; M: N=11 of 19; C: N=15 of 15) | Sometimes vigorous<br>(H: N=4 of 10; M: N=5 of 10; C: N=6 of 10) |
| **Radial Extension & Dispersion of Cerata (Stage 2)** | Coordinated, robust | Absent; no dispersion, uncoordinated/non-radial extension |
| **Response Propagation** | Whole-body; rapid; equal rates bidirectionally | Variable spread; slow; equal rates bidirectionally |

## Discussion

In characterizing the kinematics of bristling in both intact and decerebrated specimens of *Berghia stephanieae*, we have elucidated how an animal’s CNS and PNS can independently mediate different aspects of a behavior. This conclusion is borne out by the observations that 1) *Berghia*’s PNS is sufficient to effect Stage 1 bristling, in which those cerata closest to the precipitating stimulus move inward (adducting or centripetally pivoting) to sting it; and 2) the CNS is necessary to engender Stage 2 of the behavior, which involves the outward radiation of the remaining, more distal cerata to generate a dramatic, pincushion-like defensive screen around the animal’s body. Importantly, our findings extend existing work in cephalopods to challenge the notion that peripheral nervous systems are strictly limited to executing targeted movements in a subordinate manner vis-à-vis the CNS or as part of a reflex arc. To this end, and based on the mechanistic differences and similarities inherent to bristling in intact and decerebrated animals, we propose a novel two-network neuroanatomical model accounting for the kinematic sequencing observed in bristling and the underlying neural signaling driving it, in which the constituent peripheral-origin network is able to locally inhibit the corresponding central-origin network (Fig. 6).

**Figure 6.**
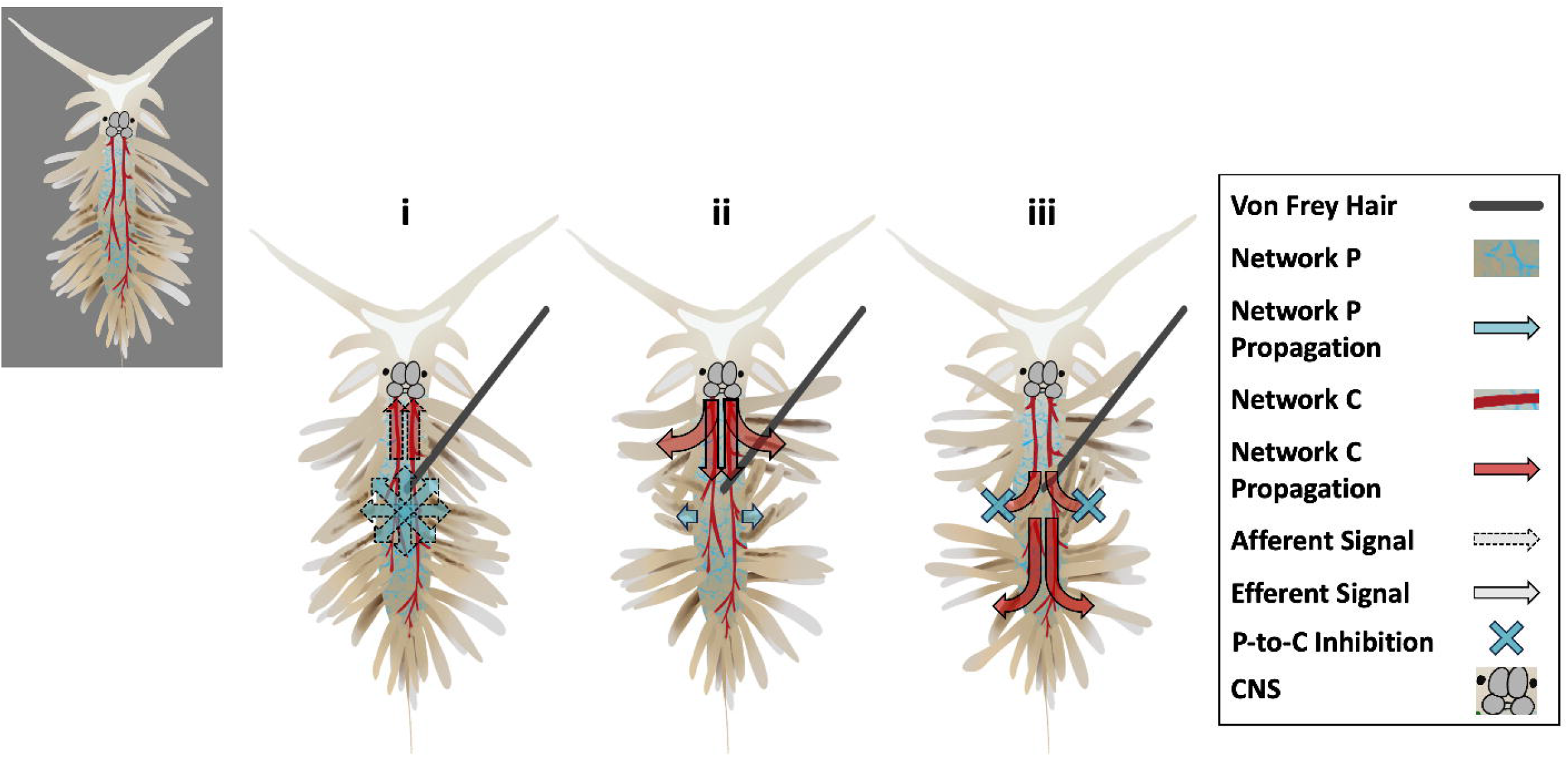
A neuroanatomical model of bristling demonstrating the kinematic sequence of the behavior and the corresponding signaling inferred to drive it. Both the retention of Stage 1 bristling and its unchanged post-stimulus onset latency following decerebration suggest that this phase of the behavior is integrated via a peripheral-origin network, Network P, that remains intact in decerebrated specimens. By contrast, the abolition of Stage 2 bristling following decerebration implicates the CNS and a centrally derived network, Network C, in generating this phase. The relative rapidity with which waves of bristling (Stage 2) propagated in intact animals suggests that Network C consists of larger-diameter processes, while Network P likely comprises a smaller-diameter neural meshwork that supports less rapid, more local signaling. In response to the midbody stimulation depicted in the figure, afferent signals are induced (Panel i) and are integrated in the periphery and CNS to give rise to efferent impulses driving ceratal movement in Networks P and C, respectively (Panel ii). Network P inhibits Network C vis-à-vis proximate cerata of the midbody to drive Stage 1 (inward) bristling in these appendages (Panel ii), while the attenuation of signals originating within the midbody portion of Network P outside of this region leaves Network C to effect Stage 2 (outward) bristling in the cerata originating elsewhere in the body (Panel ii: initial radiation of head cerata; Panel iii: complete radiation of all cerata outside Network P’s activation radius). The inset in the top-left depicts the neuroanatomical model without any corresponding signals for visual clarity. This figure was created using Procreate v. 5.3.3 for Apple iPad (Savage Interactive; Hobart, Australia).

### A neuroanatomical model of bristling

Given that the inward ceratal movement characterizing Stage 1 is apparently patterned peripherally and the fundamentally outward cerata movement observed in Stage 2 is effected by the CNS, we deduce that, at a minimum, the neural infrastructure mediating bristling comprises two distinct neural networks. Both the retention of Stage 1 bristling in decerebrated *Berghia* and the equivalent post-stimulus latencies to initial Stage 1 induction between intact and decerebrated specimens suggest that this phase of the behavior is driven by a peripherally situated efferent network, Network P (cyan neuropil in Fig. 6). By contrast, the elimination of Stage 2 bristling in decerebrated specimens implicates a centrally derived network, Network C (crimson neuropil in Fig. 6), in conveying output that drives the coordinated outward radiation of cerata sufficiently distal to the incident stimulation. The two-network framework is furthermore supported by the observation that when the midbody is stimulated, efferent information is relayed from the CNS to the caudal body to induce Stage 2 bristling in the latter region and remains unimpeded during its transit through the midbody, where the concurrent transmission of outputs driving Stage 1 bristling in that region is occurring. Finally, the presence of incoherent, spontaneous ceratal movement in decerebrated specimens suggests that Network C normally supplies tonic inhibition to those motor neuron pools mediating ceratal movement, and that during bristling, this tonic inhibition is lifted or otherwise surmounted by excitatory input.

Each of these two bristling networks may associate with dedicated sensory neuron populations, conducting parallel streams of afferent information to sites of peripheral or central integration; alternatively, a single pool of sensory neurons may carry the inputs ultimately eliciting both stages of bristling by projecting divergently to integrative elements in the PNS and CNS. In either case, the centripetal pivoting of all cerata within a fixed radius of the impinging stimulus to sting it during Stage 1 bristling suggests a generally radial organization of the afferent projections associated with Network P, with Network-C-associated afferents conveying sensory traffic directly to the CNS (Fig. 6, Panel i).

That the waves of uncoordinated ceratal recruitment in decerebrated specimens traveled significantly more slowly and, on average, over shorter distances along the body than those generated in intact animals implies that Network P, which would remain the default signaling framework in decerebrated animals, consists of smaller-diameter neurites marked by slower conduction velocities relative to the larger-diameter tracts presumed to comprise Network C, which accommodates the more rapid, whole-body signal propagation inherent to Stage 2 bristling in intact animals. Based on anti-tubulin staining conducted in juvenile *Berghia*, including near the ceratal bases (C. Tait, personal communication), and a variety of other anatomical stains pursued in both sensory and non-sensory organs of other gastropods [38-45], we further infer that Network P is organized as a highly ramified cutaneous and/or subepithelial nerve plexus spanning *Berghia*’s body. The somata contributing to this network are possibly situated within the plexus itself, as in many gastropod species in which peripheral neuroanatomy has been characterized [38-45]; alternatively, they may be housed in as-yet-unidentified ceratal ganglia analogous to those identified within the considerably larger and less numerous cerata of *Melibe* [36]. By contrast, we deduce that the neuronal somata giving rise to the processes comprising Network C originate in central ganglia and include elements homologous to those neurons implicated in directing more general ceratal movement in *Hermissenda* and *Phestilla* [32, 37]. The ramifying of the associated neurites may mirror those of pleural nerves in *Melibe*, branches of which ultimately innervate ceratal ganglia in that animal [36], with these processes likely facilitating rapid conduction and diverging to produce large innervation fields, analogous to the large, central motor neurons in the pteropod sea slug *Clione limacina* that innervate that animal’s parapodia to mediate swimming [46]. Indeed, these features of Network C would be sufficient to account for the statistical simultaneity of Stage 2 bristling induction in the head and midbody regions when the caudal locus was stimulated, on the assumption that afferent signals originating in the caudal body, when integrated in the CNS, give rise to motor outputs that exhibit equally rapid transit times to the head and midbody cerata due to the specific branching patterns of Network C in those regions.

How is it that two distinct neural networks can drive different modes of movement in two different subsets of cerata? We posit that this duality involves circuitry wherein Network P is able to locally and unidirectionally inhibit Network C. In this configuration, the midbody stimulation portrayed in Fig. 6, for example, would induce efferent signals in both networks (Panel ii), with the relative signaling intensity of Network P in the midbody region sufficient to inhibit Network C activity, promoting the effect of Network P, Stage 1 bristling, in the stimulus-proximate midbody cerata (Panels ii and iii); notably, this inhibition in the midbody occurs without interfering with the transit of efferent signals from the CNS to the caudal region via Network C. In the head and caudal regions, as well as areas of midbody sufficiently distal to the stimulus, the intensity of Network P signaling would have attenuated at sufficient distances from the site of induction to allow Network C to dominate in these body regions by default, eliciting Stage 2 bristling in all cerata outside of Network P’s activation radius (Panel iii). At a circuit level, this mechanism would presuppose two distinct pools of peripheral primary motor neurons promoting unique modes of ceratal movement—one directing inward, retaliatory action (Stage 1) and associated with Network P and another driving outward, preemptive ceratal radiation (Stage 2) and associated with Network C—in which Network P would have the capacity to inhibit Network C either at or upstream of the motor neuron layer. While this sort of unidirectional inhibitory synaptic architecture has precedent in promoting the singleness of behavioral output within the CNS, both in gastropods [22, 47, 48], our model entails a novel hierarchical relationship between the PNS and CNS that biases one stage of the bristling behavior over the other as a function of stimulus proximity. Anatomically, the interaction between the bristling-mediating networks in *Berghia* might on the one hand occur within overlapping subepithelial neuropil or at somata distributed at neural branch points, as has been observed between central and peripheral elements in other gastropods [40, 42, 49]. Alternatively, this complexing could occur in hypothetical ceratal ganglia within the body, recalling the general interaction between central and peripheral neurons posited in the peripherally situated tentacle and rhinophore ganglia of the gastropod *Pleurobranchaea californica* [50] or within the gill and osphradial ganglia of *Aplysia* in the context of the gill withdrawal [49].

### Interactive, distributed patterning of non-reflexive behavior across the central and peripheral nervous systems

Despite the stereotypy inherent to certain aspects of bristling and the difficulty in distinguishing voluntary from involuntary action in gastropods, for whom the concept of volition is poorly defined, the behavior deviates appreciably from the criteria that classically define a reflex [51]. Although both vertebrate and invertebrate reflexes, such as scratching [52, 53] or startling [54], may involve multiple constituent movements and, in the case of scratching, may also be directed at the stimulus precipitating it, bristling entails a sequence of two disparate modes of movement within the same set of appendages, in which individual cerata can flexibly reallocate their roles from one stage of bristling to the other across successive defensive episodes depending on their proximity to the stimulus; this contrasts bristling with the synergistic limb and body movements that constitute scratching and reflex action more generally. This lack of synergy is recapitulated at the level of the circuitry we infer to mediate bristling: whereas the CNS and PNS integrate reflexes cooperatively along sensorimotor arcs, bristling is deduced to involve an antagonistic interaction between these two systems based on the suppression of Stage 2 bristling, an otherwise whole-body response, in the body region being stimulated. Moreover, the directed portion of bristling, Stage 1, can be mediated exclusively in the periphery, whereas targeted reflex movement depends upon central integration and pattern generation [53].

Bristling, to our knowledge, thus represents not only the first instance in which the brain and PNS autonomously pattern different phases of a single animal behavior, but one that also features a novel hierarchical dominance mechanism in which the PNS can override the CNS. Although the interaction of two or more neural networks in driving behavior is not unique to bristling in *Berghia,* neither does it precisely mirror similar neuroethological paradigms identified in other animal taxa. In several cnidarians, such as the jellyfish *Aglantha digitale* and *Carybdea marsupialis* and the freshwater polyp *Hydra vulgaris*, multiple functionally discrete (though sometimes anatomically overlapping) conduction systems subserve individual behaviors, such as distinct modes of swimming, elongation and contraction, defensive crumpling, and light-induced locomotion [55-58]. The sequencing intrinsic to somersaulting behavior in *Hydra* has been speculated to involve reciprocal inhibition between two distinct populations of neurons, distinguishing it from the unidirectional inhibition we infer to underlie the *Berghia* bristling sequence [59]. Furthermore, the neuronal populations implicated in *Hydra* somersaulting are thought to belong to the same ectodermal network, whereas there is evidence that the PNS and CNS in gastropods, and hence the peripheral- and central-origin networks we deduce mediate bristling, may be embryologically distinct based on when their ostensible precursors differentiate during development [60].

Among cephalized animals, the octopus represents a singularly robust model for studying the interaction between nervous systems. Hochner [5] has proposed an “embodied intelligence” framework to account for how the brain and PNS, distributed across the eight arms in these animals, interact to generate behavior. In this organizational scheme, the periphery does not merely execute individual motor commands issued by the brain; rather, interactions between the sensory infrastructure of the arms, their dynamic physical constitutions (reflecting the arms’ effectively infinite degrees of freedom), and the brain interface synergistically to pattern movements. While the embodied intelligence framework ascribed to the octopus could conceivably account for behavior involving more structurally complex, behaviorally versatile, and physically flexible appendages in gastropods, such as tentacles, the different networks apparently driving bristling in *Berghia* appear to interact antagonistically via peripheral-to-central inhibition, rather than synergistically, to evoke different phases of that behavior. Nevertheless, the peripheral-central division of neural labor paradigm we deduce to generate bristling is not so dependent on *Berghia*’s specific neuroanatomy or body plan that it would be unreasonable to expect to identify it in other behaviors across animal phylogenies.

### Future directions

The findings we detail here further challenge traditional conceptions in the field of the PNS’ capacity to shape complex behavior independently of the brain and, as such, stand to motivate an interest in utilizing state-of-the-art molecular, optogenetic, and computational tools to further interrogate the physiology of this largely overlooked division of the nervous system. With respect to bristling itself, significant insights could be yielded from the generation of transgenic *Berghia* expressing genetically encoded calcium or voltage indicators (GECIs/GEVIs) in both central and peripheral neuronal populations, an effort currently being pursued in this species. Even prior to this breakthrough, efforts in the immediate future can focus on advancing our understanding of bristling’s central neuronal correlates, given our present ability to load fast absorbance voltage-sensitive dyes into the *Berghia* CNS: this imaging approach has already afforded us single-neuron, single-spike resolution across significant portions of the animal’s isolated brain [61]. Immobilized whole-animal or reduced preparations retaining some portion of the mantle and associated cerata could furthermore be appropriately stimulated to elicit bristling, allowing for the direct observation of population-level neuronal dynamics over the full course of the behavior’s underlying sensorimotor integration; already, we are able to record from and stimulate individual neurons in the CNS of intact animals using sharp electrodes (see Fig. 1C, inset), which enables the study of synaptic relationships between bristling-related neurons identified in the optical records. Ultimately, the ability to image and stimulate neurons in transgenic *Berghia*, and in particular *Berghia* juveniles, whose mantles retain degrees of transparency during stages of ceratal development [34] and would therefore be especially amenable to whole-body imaging, could facilitate the evaluation of our bristling model by enabling the visualization of signal propagation associated with the behavior within those networks hypothesized to mediate it. The mechanistic principles gleaned through such efforts could, at best, advance a more general understanding of how the brain and PNS interact to generate concerted behavior and would epitomize the unique potential represented by the gastropod model system to the field of neuroscience.

## Acknowledgments

We wish to thank Viral Mistry and Babita Thadari for their experimental assistance, Dr. Evan Hill for working on an early version of Figure 6 and for his comments on the manuscript, Dr. Cheyenne Tait for her correspondence and sharing of unpublished histological data, and our *Berghia* Brain Project collaborators for introducing us to the bristling behavior and for their insights and feedback throughout the course of the project. This work was supported by NIH grants U01NS108637 and R01NS121220.

## Materials and Methods

### Animals

Adult specimens of *Berghia stephanieae* ranging in length from 6 to 21.5 mm were sourced from the commercial supplier ReefTown.com (Boynton Beach, FL) or obtained from the research laboratory of Dr. Paul S. Katz at the University of Massachusetts at Amherst; additional *Berghia* were drawn from a self-sustaining, homegrown population deriving from animals obtained from our two suppliers and raised in a 20-gallon marine system. Specimens used in the study were maintained in aerated, approximately 1-liter bowls of artificial seawater (ASW; Instant Ocean, Blacksburg, VA) at a specific gravity of 1.024-1.025 and maintained between 24-26°C using a heating pad to approximate the average temperature of the animals’ native habitat. Animals were kept under ambient light-dark conditions, with all experiments conducted during the daytime. Specimens were fed *Exaiptasia diaphana* (obtained as *Aiptasia* from Carolina Biological Supply, Burlington, NC) once every three days and were included in experiments approximately 24 hours after feeding. Animals were transferred from their bowls to experimental enclosures using a large-diameter plastic Pasteur pipette. As this transfer process periodically induced ceratal autotomy, a phenomenon documented in aeolids when individual appendages endure sustained pressure [30, 31], we only utilized *Berghia* that were judged to have retained a large majority (at least 75%) of their cerata. The body lengths of all specimens used in the study were measured prior to experimentation.

### Behavioral Procedures

To facilitate maximum spatiotemporal resolution, bristling was primarily studied in immobilized *Berghia*. Although specimens sometimes exhibited brief, spontaneous episodes of modest (i.e., non-bristling-like) ceratal elevation and/or engaged in vigorous attempts to break free of the vacuum suction in the initial period following the introduction of immobilization, these behaviors typically subsided over the 5-10 minutes after the vacuum was initialized.

The head, midbody, and caudal stimulation regions (Fig. S1B) were stimulated in a random order with a 3.61-gauge VFH monofilament, which delivered a calibrated force of 3.9 mN to the mantle. Each locus was stimulated 2-4 times in two-minute interstimulus intervals; for every animal, the replicate video featuring the most accurate targeting at each stimulation site was selected for kinematic analysis. Immobilization was achieved through the use of a customized vacuum apparatus consisting of a 5.5-cm plastic Petri dish with three 1.25-mm holes drilled in a row into the bottom-center of the dish and spanning 6.0 mm; these holes facilitated gentle vacuum suction from below (Fig. S1A). An appropriately sized wedge of Whatman Grade-1 filter paper (Sigma-Aldrich, St. Louis, MO) was cut, wetted, and placed directly over the central holes of the chamber to facilitate suction of the animals. Suction was initialized to immobilize the animal by vacuuming its foot to the filter paper, precluding forward locomotion, turning, and contraction without distorting the animals’ body or noticeably impeding bristling. ASW perfusion into the Petri dish via a Mariotte’s bottle was then balanced relative to vacuum suction to ensure a steady-state water level in the dish. Specimens were given 5-10 minutes to adjust to vacuum immobilization before the beginning of stimulation, with the commencement of experimentation contingent on both the cessation of brief, vacuum-evoked episodes of ceratal elevation and vigorous attempts to break free of the suction. Any animals failing to acclimate over this period were not used in the study. In the event that specimens escaped from vacuum suction between trials, suction was reinitialized, and specimens were allowed 5-10 minutes to reacclimate before stimulation resumed; data collection was terminated in any animal that escaped vacuum suction three times.

Pilot experiments attempting to record freely locomoting animals responding to tactile stimulation through a stereoscope revealed that such videos could not be reliably obtained due to the rapid locomotion of animals to areas outside of the microscope’s field of view; the specimens’ locomotion likewise precluded precise tactile stimulation with a VFH. Freely locomoting animals were therefore recorded with an overhead camera (see the “Video Acquisition” subsection), and focal application of 1M NaCl was used to elicit bristling in freely locomoting specimens, with stimulations limited to the head and caudal regions. This concentration of NaCl, which possesses nearly twice the osmolarity of normal seawater and was noxious to *Berghia*, drove the same qualitative bristling response as generated by VFH application and without prompting specimens to drop cerata (which was observed in pilot experiments featuring 2M and more concentrated NaCl solutions); by contrast, ASW equivalently applied during pilot studies did not typically provoke bristling or accompanying defense behaviors.

Freely locomoting *Berghia* were individually placed in an 85-mm glass Petri dish (hereafter referred to as the “arena”) filled with room-temperature (22-23°C) ASW and allowed to adapt to the environment for five minutes before experimentation commenced. A 23-gauge needle mounted on a 1-mL syringe was used to apply 20µL of 1M NaCl to either the head or caudal region over the course of approximately 1.5 seconds. Applications were repeated six times at one anatomical site, with two minutes separating applications; replications featuring imprecise targeting or specimens crawling along the edges of the arena were discarded. Animals were then transferred to a holding dish, during which time the ASW in the arena was changed. After specimens were given five minutes to reacclimate to the arena, the other anatomical site was similarly stimulated; the order of stimulation (head-caudal or caudal-head) was randomly determined for every specimen. Because *Berghia* frequently displayed thigmotaxis, individuals observed crawling vertically around the edges of the arena, from which full-body video could not be reliably acquired, were gently repositioned to the center of the arena between trials using plastic forceps.

### Decerebration Protocol

Specimens to be decerebrated were first anesthetized by placing them in an elastomer-lined dish filled with a 350 mM solution of MgCl_2_ for 10-15 minutes. Anesthetized animals were then pinned down, and an approximately 1-mm-long dorsal incision was made superficial to the CNS to facilitate its gentle excision using forceps and fine scissors. Decerebrated specimens were placed in separate, aerated holding bowls for 24 hours, during which time the incision invariably fused. These specimens were then run through the same tactile stimulation paradigm under immobilization employed with intact animals.

### Video Acquisition

All experimental trials were filmed for later kinematic analysis. Image stacks acquired of immobilized animals through the ORCA-Flash4.0 camera were collected in the associated software, HD Image (v. 4.6.1.2), and subsequently compiled into videos using Fiji (v. 2.1.0/1.53c; [62]). In the free-locomotion paradigm, an AmScope MU300 color CMOS camera (Irvine, CA) was used with an SMC Pentax-M 28-mm F2.8 lens (Lewisville, TX) and was suspended approximately 0.75 m above the testing arena to capture the full arena during recording. Video was acquired using AmScope software for Windows 10 (v. 4.11.18573) at 25 frames per second (fps) and at a resolution 768 x 1024 pixels.

### Data Analysis

All experimental videos were reviewed using Fiji, in which kinematic measurements involving distances, angles, and times were made; distance measurements initially made in pixels within Fiji were ultimately converted to metric units based on pre-experimental measurements of specimens’ body lengths. Based on the early observation that many cerata fanned out from one another during bristling, we developed a method for quantifying the extent of this dispersion. First, the coordinates corresponding to individual cerata, as marked by the position of the cnidosac at their tips, were manually registered in every video featuring immobilized animals across two specific frames: 1) the one immediately preceding stimulation (control) and 2) the one in which the cerata were judged to have maximally dispersed (bristling-maximal). Custom Matlab code (MathWorks, Natick, MA) was then utilized to calculate the distance between every ceras and its closest neighbor in both the control and bristling-maximal frames, after which the mean interceratal distance 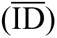 was calculated at each frame by averaging unique, pairwise distances between cerata. In those trials in which ceratal dispersion increased over the course of bristling, a further metric, the “bristling magnitude” (|B|), or normalized change in ceratal dispersion, was computed as

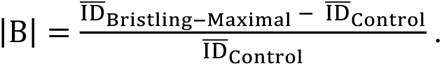

To calculate the post-stimulus ceratal response latencies in the same set of videos, we reviewed videos in a frame-by-frame manner. The latency to bristling induction in a given body region was judged based on the first post-stimulus pivot of a ceras in that region, while the latency to whole-body bristling, when observed, was determined around the first frame in which cerata across all body regions had begun pivoting. To calculate the speed at which bristling propagated from one body locus to another, distances between two loci were measured in Fiji and divided by the associated difference in bristling induction latency.

To compute the crawling speeds of freely locomoting animals, we utilized the Fiji plugin AnimalTracker [63] to derive trajectories for the centroid of each specimen; we then implemented custom Matlab code to calculate instantaneous speeds on every video frame, binned in nonoverlapping 0.5-s increments. Any artifactual measurements in speed resulting from the body contraction that rapidly followed stimulation were replaced with values estimated through linear interpolation of speeds in adjacent time bins.

All statistics were computed using GraphPad Prism (v. 9.4.0, San Diego, CA), R [64], and Microsoft Excel for Mac (v. 16.63.1, Redmond, WA). As all data sets were either normally or log-normally distributed, we used parametric testing to ascertain statistical significance at a threshold of *p*<0.05 (two-tailed), with Shapiro-Wilk tests utilized to assess normality and both Brown-Forsythe and Bartlett’s tests employed to judge homogeneity of variance. Where necessary, data were log-transformed to render them normal and/or homoscedastic; in these cases, we reported metrics associated with statistical models (e.g., *F* and *p*) based on log-transformed data while presenting descriptive statistics relative to raw data. Student *t*-tests and one- or two-way analyses of variance (ANOVAs) were used to compare means across measurements, with Tukey’s post-hoc tests correcting for multiple comparisons accompanying the latter analyses as appropriate. Paired *t*-tests or repeated-measures (RM) ANOVAs were employed to analyze metrics that changed within individual specimens over time. For freely locomoting animals, in which measurements were replicated in individuals, weighted-means *t*-tests were computed using the R package “weights” [65] to compare certain measurements as functions of head and caudal stimulation, with simple means derived from individual animals weighted with the reciprocals of their corresponding standard errors (i.e., 1/SEM); the aggregate means and SEMs across animals for these measurements are reported in the Results section and similarly reflect this weighting. We used an RM-ANOVA-like mixed-effects model with the Geisser-Greenhouse correction to detect significant changes in the forward crawling speeds of *Berghia* prior to and following stimulation; this model utilized aggregate mean speeds reflecting the same weighting method as above at each time step for every animal.

### Supplemental Information

**Figure S1.**
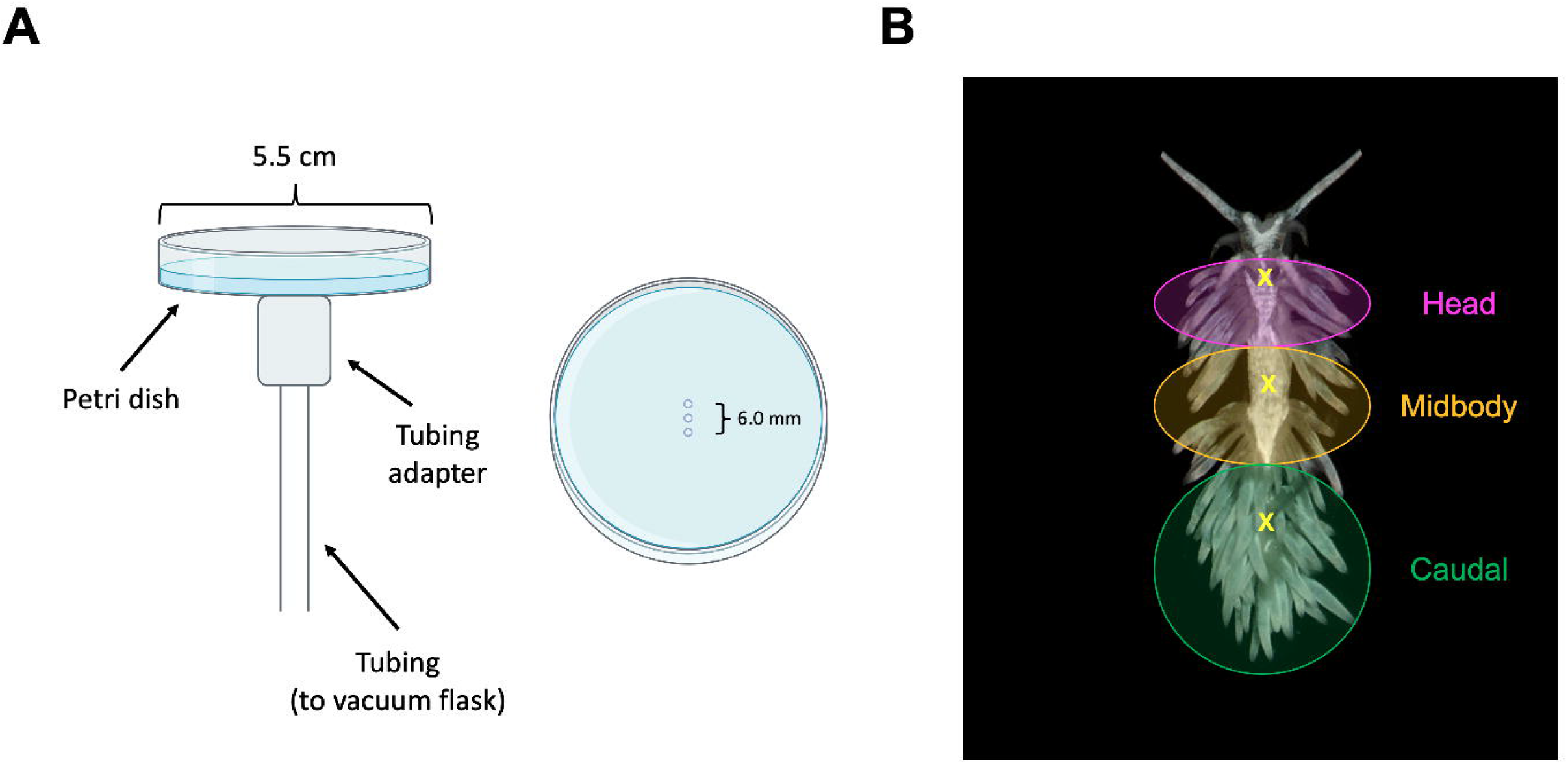
Methodology for tactile stimulation. (A) To facilitate precise stimulation and high-resolution recording, a customized vacuum chamber was utilized to immobilize animals against a piece of filter paper (not shown) against the chamber floor with gentle suction. Shown are lengthwise (left) and overhead (right) views of the apparatus; the three holes mediating suction are visible in the overhead view and are not drawn to scale. Petri dish illustrations were sourced from BioRender.com. (B) Immobilized specimens were stimulated with a von Frey hair at three dorsomedial body loci (each of which is indicated by an “X”): the head, midbody, and caudal sites. For subsequent analysis, all cerata were grouped into one of three regions (colored circles) according to the proximity of their insertion points to the three stimulation loci. Freely locomoting animals were stimulated with 1M NaCl in only the head and caudal regions.

**Figure S2.**
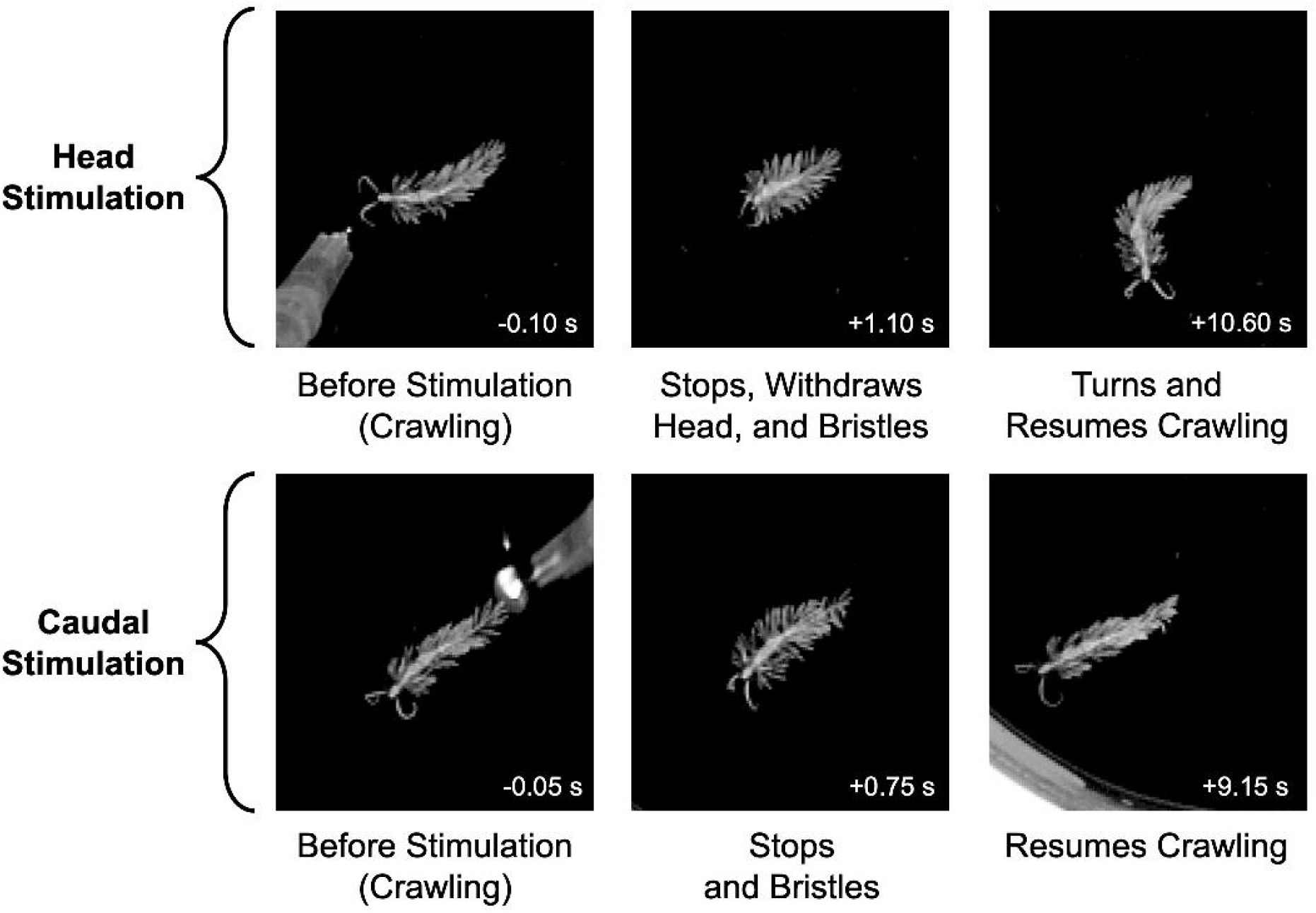
Aversive dorsal stimulation of freely locomoting *Berghia* elicited a suite of defensive behaviors, including bristling. Application of 20 µL of 1M NaCl over the course of 1.5 s to *Berghia’*s head and caudal regions triggered bristling and a concurrent cessation of forward locomotion in this pair of representative behavioral sequences. Head stimulation usually elicited turning, while caudally stimulated animals typically retained their previous headings (see Fig. S3). The resumption of forward crawling typically coincided with the initiation of ceratal relaxation and, to whatever extent aversive stimulation drove body contraction, a reversion of the body to its resting length. Time indices in each panel are reported in relation to stimulation. See also Videos S2 and S3.

**Figure S3.**
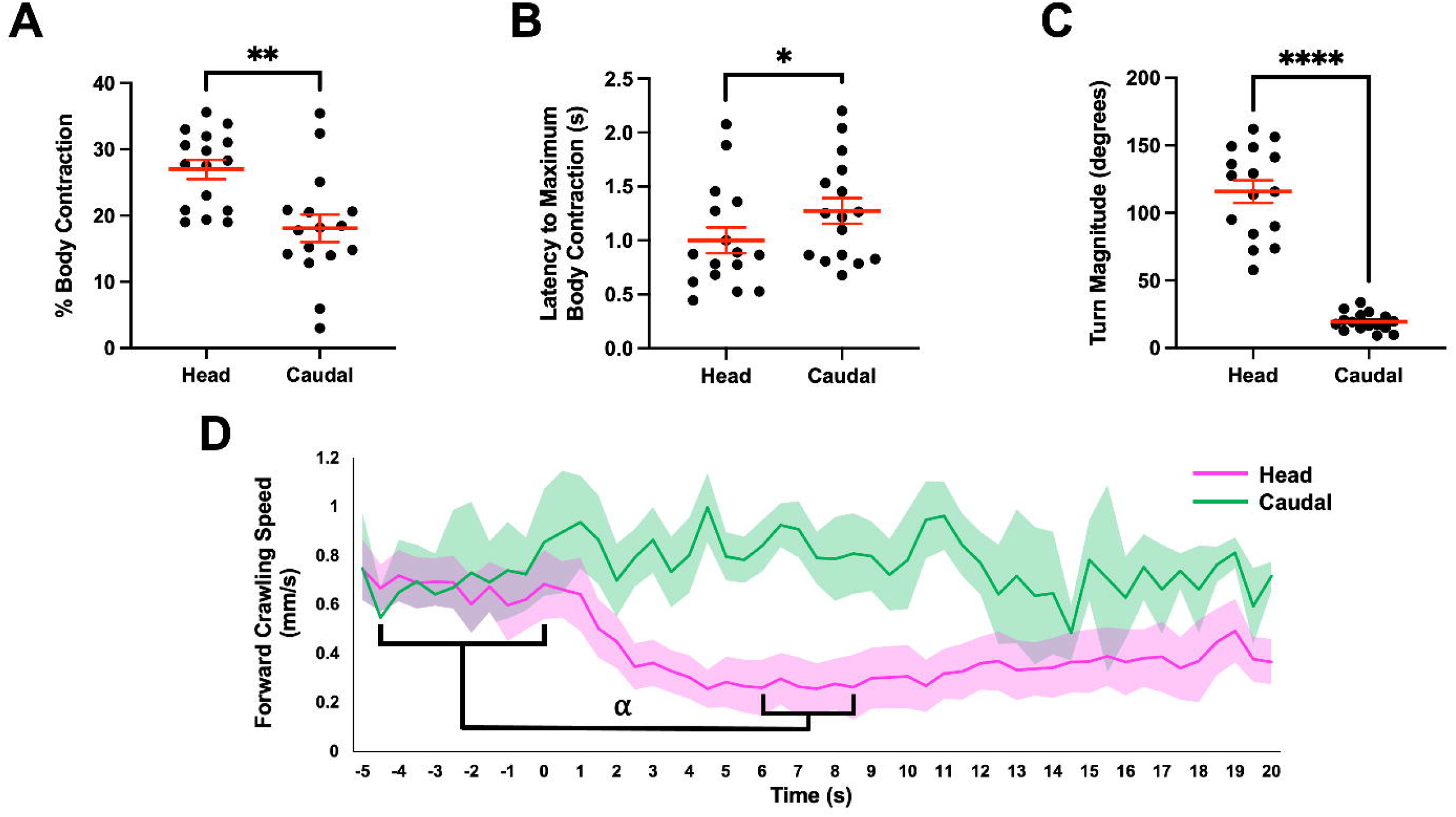
Head stimulation drove more dramatic accompanying defensive behaviors in freely locomoting animals than tail stimulation. Stimulating the head of *Berghia* (N=16) with 20 µL of 1M NaCl effected (A) more extensive, (B) more rapid body contractions and (C) larger-magnitude turns than caudal stimulation [weighted-means *t*-tests: (A): *t*(25.63)=3.34, *p*=0.0026; (B): *t*(29.86)=-2.45*, p*=0.021; (C): *t*(16.31)=12.04*, p*<0.0001]. The onset and termination of turning were judged based on the initial bending and relinearization of the body, respectively. As caudal stimulation typically evoked negligible turning, the corresponding temporal measurements could not be reliably made in these trials due to the subtlety and rapidity of body bending. Despite the lack of a solid stimulus, both stages of bristling characterized in the tactile stimulation paradigm were observed in response to punctate application of NaCl, with those cerata participating in Stage 1 adducting toward the dorsal application site. The displayed statistics in Panels A-C reflect weighted means (with simple means from individual specimens weighted with the reciprocal of the associated SEMs), while the plots themselves reflect raw data distributions. ****, *p*<0.0001; **, *p*<0.01; *, *p*<0.05. (D) A repeated-measures-ANOVA-like mixed-effects model with the Geisser-Greenhouse correction revealed that head stimulation but not caudal stimulation significantly slowed forward crawling. We looked for significant changes in the average speeds across specimens over the pre- and post-stimulatory epochs, respectively defined as the 5 s preceding and 20 s following stimulation, resulting from head and caudal stimulation. For head stimulations, analysis revealed significant variations in speed [*F*(6.71,99.62)=3.99, *p*=0.0008], with a significant deceleration of over 50% on average between the pre-stimulatory epoch (-4.5 to 0 s) and the period between 6 and 8.5 s following stimulation; by contrast, the same analysis detected no significant changes in speed resulting from caudal stimulation [*F*(4.80,51.40)=2.28, *p*=0.0631]. Post-head-stimulus deceleration was consistent with the need to accommodate the typically accompanying head withdrawal, as well as the observation that animals so stimulated executed large-magnitude turns; forward crawling at nominal speeds would be expected to impede the efficiency of these larger, more sustained rotations. The more pronounced effect of head relative to caudal stimulation on crawling speed, elicited turn magnitude, and contractile magnitude and speed collectively intimated a greater urgency in animals’ responses to head stimulation, despite the lack of significant head vs. caudal differences in bristling magnitude and characterized bristling latencies (see Figs. S4 and S5). The displayed statistics reflect those Tukey post-hoc tests that indicated significant pairwise comparisons across the measurement window (see Table S1); ⍺, 0.0076<*p*<0.05. Data were binned in 0.5-s increments, with trend lines and error bands respectively derived from aggregate means and SEMs reflecting the above weighting scheme.

**Figure S4.**
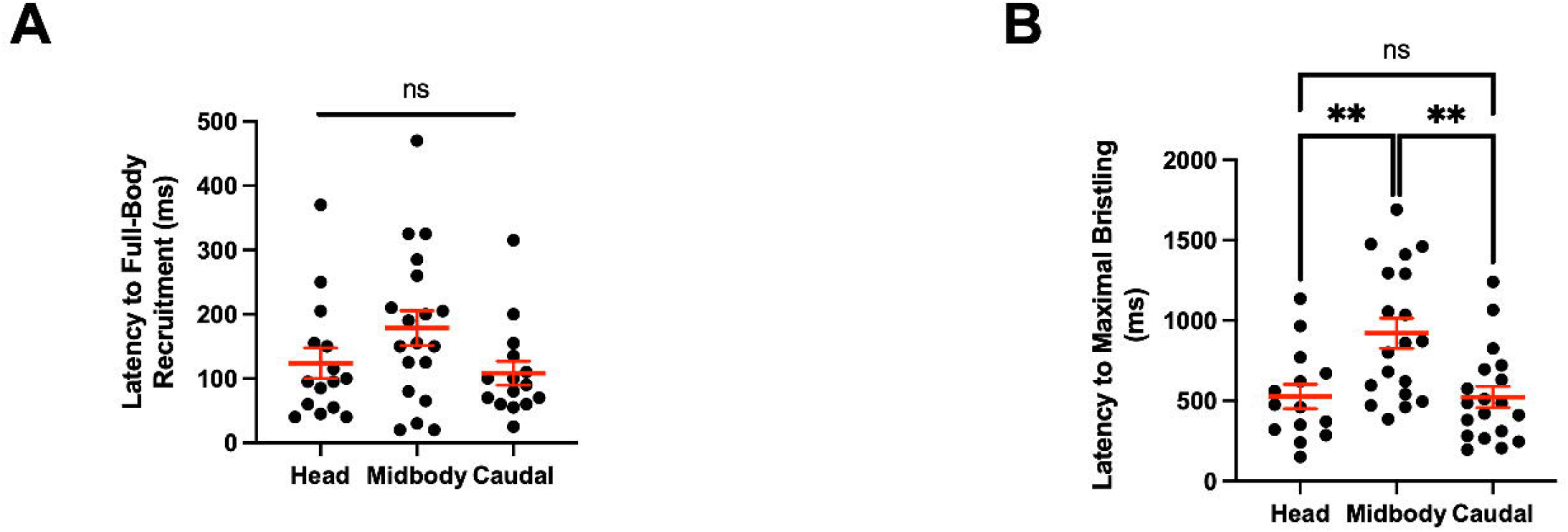
Post-stimulus latencies to full-body bristling were invariant across stimulus regions, while latencies to maximal bristling magnitude were longer following midbody stimulation. (A) There were no significant differences in the post-stimulus latencies to full-body ceratal recruitment (i.e., the time required for all cerata to respond to stimulation) as a function of stimulation site [one-way ANOVA, *F*(2,46)=1.13, *p*=0.331]. (B) By contrast, the average latency to the time at which maximal bristling magnitude (see Fig. S5) was achieved was significantly greater when the midbody was stimulated than the head or caudal regions [*F*(2,49)=8.11, *p*=0.0009; Tukey’s tests: *p*=0.0053 for head vs. midbody, *p*=0.0022 for midbody vs. caudal, *p*=0.9998 and head vs. caudal]. This could conceivably reflect less sensitivity to noxious stimuli in the midbody region, slower afferent transmission therein, or a reduced urgency to bristle when this area was stimulated. **, *p*<0.01.

**Figure S5.**
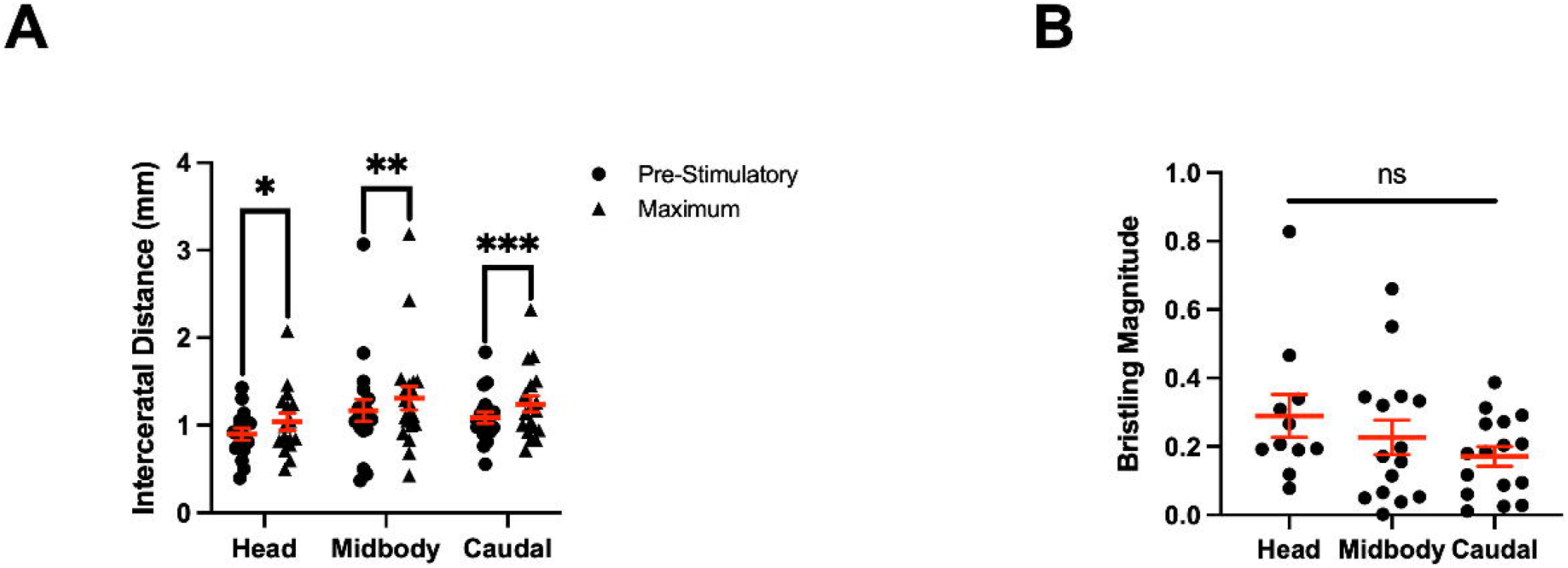
Bristling magnitude did not significantly vary across stimulated body loci. (A) The distance between those cerata participating in Stage 2 bristling, as measured between the cnidosacs (i.e., ceratal tips) before stimulation and at the fullest extent of bristling, increased significantly over the course of the behavior (i.e., between the time immediately preceding stimulation to the time when bristling was judged to have achieved its maximum extent) in the majority of trials involving head (N=11 of 17), midbody (N=15 of 20), and caudal stimulation (N=16 of 20) in immobilized animals [paired *t*-tests: head, *t*(15)=2.570, *p*=0.0213; midbody, *t*(19)=3.241, *p*=0.0043; caudal, *t*(18)=4.641, *p*=0.0002]. (B) Bristling magnitude, calculated as the pre-stimulatory-normalized change in Stage 2 ceratal dispersion, did not significantly vary as a function of stimulus site [one-way ANOVA, *F*(2,38)=0.7911, *p*=0.4607]. ***, *p*<0.001; **, *p*<0.01; *, *p*<0.05; ns, not significant.

**Table S1.** A table of Tukey’s post-hoc tests revealing significant pairwise differences in forward crawling speed between select pre- and post-stimulatory times (binned at 0.5 s) when 20 µL of 1M NaCl was applied to the head region of freely locomoting *Berghia* (N=16). All other Tukey’s tests computed for crawling speeds measured over the period spanning 5 s before (-5 s) and 20 s following stimulation did not meet the multiplicity-adjusted threshold for significance. These tests were run following the revelation of overall significance with a repeated-measures-ANOVA-like mixed-effects model with the Geisser-Greenhouse correction.

| Times Compared<br>(s, 0.5-s bins) | Adjusted $p$ value |
| --- | --- |
| -4.5 vs. 8.5 | 0.0259 |
| -3.5 vs. 6 | 0.0282 |
| -3.5 vs. 7.5 | 0.0476 |
| -3.5 vs. 8.5 | 0.0186 |
| -3 vs. 6 | 0.0314 |
| -2.5 vs. 6 | 0.0278 |
| -2.5 vs. 8.5 | 0.0102 |
| -2 vs. 7.5 | 0.0144 |
| -2 vs. 8.5 | 0.0162 |
| -1.5 vs. 6 | 0.0198 |
| 0 vs. 8.5 | 0.0234 |
| -2 vs. 6 | 0.0076 |

**Video S1. Bristling elicited in *Berghia stephanieae* in response to an aversive tactile stimulus.** This video plays in real time.

**Video S2. A freely locomoting *Berghia* responding to the application of 20 µL of 1 M NaCl to the head region.** In response to stimulation, this specimen bristled, while concurrently contracting its body and withdrawing its head. It subsequently pursued a large-magnitude turn away from the stimulus, during which time its cerata and head relaxed and its equilibrium body length was restored. This video and Video S3 are accelerated to 3X real time.

**Video S3. A freely locomoting *Berghia* responding to the application of 20 µL of 1 M NaCl to the caudal region.** As with the animal in Video S2, this specimen bristled, while concurrently contracting its body and withdrawing its head, though not as dramatically as its head-stimulated counterpart. The animal continued along a largely undeviating trajectory while its defensive maneuvering subsided.

**Video S4. A vacuum-immobilized *Berghia* responding to von Frey hair stimulation of the head region.** In response to stimulation (t=0 ms), many cerata in the head region adducted and/or centripetally pivoted to envelop the tip of the VFH (Stage 1 bristling), while more posterior head cerata and those originating in the midbody and caudal regions dispersed above and lateral to the mantle while pivoting anteriorly (Stage 2 bristling). This video is slowed to 1/20^th^ real time, as are Videos S5-S7. Animals in Videos S5-S8 are similarly immobilized.

**Video S5. *Berghia* responding to von Frey hair stimulation of the midbody region.** Stimulation drove nearly all midbody cerata and some posterior head cerata to adduct and/or centripetally pivot to sting the VFH (Stage 1), while more anterior head cerata and those in the caudal region dispersed radially (Stage 2).

**Video S6. *Berghia* responding to von Frey hair stimulation of the caudal region.** The impinging VFH elicited adduction and centripetal pivoting of proximate caudal cerata towards it (Stage 1), while more anterior caudal cerata and all of those originating in the head and midbody regions pivoted anteriorly and either superior or lateral to the body (Stage 2). A minority of posteriorly situated caudal cerata exhibited minimal response to stimulation.

**Video S7. A decerebrated *Berghia* specimen responding to von Frey hair stimulation of the head region.** VFH stimulation evoked both adduction and centripetal pivoting in some of the nearby cerata (Stage 1). While movement was also driven in more distal cerata, this movement was unpredictable and uncoordinated with respect to both speed and direction across cerata, mirroring the spontaneous ceratal activity preceding the stimulus. Stage 2 bristling was not observed in this specimen.

**Video S8. A decerebrated *Berghia* specimen exhibiting spontaneous ceratal movement.** Ceratal movement in decerebrated animals was persistent and uncoordinated absent stimulation, possibly reflecting disinhibition owing to the removal of the CNS. This video plays in real time.

